# Circulating cell-free RNA reflects inflammatory and airway remodeling signatures in an equine model of asthma

**DOI:** 10.1101/2025.10.25.684590

**Authors:** Birong Zhang, Hanna Agdour, Miia Riihimäki, Sanni Hansen, Amanda Raine

**Affiliations:** Department of Medical Sciences, Uppsala University; Science for Life Laboratory; Department of Clinical Sciences, Swedish University of Agricultural Sciences; Department of Veterinary Clinical Sciences, University of Copenhagen

**Author notes:** Equal contribution (Birong Zhang, Hanna Agdour).

## Abstract

Circulating cell-free RNA (cfRNA) offers a minimally invasive means to monitor dynamic gene expression across tissues, providing a promising source of biomarkers for inflammatory diseases. Equine asthma, a prevalent respiratory disorder, represents a valuable translational model for the human condition. Here, we present the first transcriptomic analysis of plasma cfRNA from asthmatic and healthy horses, alongside comparative profiling of bronchoalveolar lavage and whole blood. Tissue-of-origin deconvolution revealed asthma-associated immune cell redistribution, characterized by reduced B-cell and increased T-cell signatures that were undetectable in the conventional sampling. Differential expression analysis identified up-regulation of key alarmins and transcripts linked to airway remodeling and immune activation. Cross-compartment correlation demonstrated that cfRNA captures distinct yet biologically relevant molecular signatures consistent with asthma-related pathways. These results establish cfRNA as a sensitive complementary approach for characterizing molecular perturbations in asthma pathogenesis and highlight its potential for biomarker discovery across inflammatory and allergic airway diseases.

## Introduction

Equine asthma (EA) is a naturally occurring chronic inflammatory airway disease that shares clinical and molecular features with human asthma. Bronchoalveolar lavage (BAL) cytology remains the diagnostic gold standard in EA but provides limited information on immune and tissue-level processes. Whole-blood transcriptomes can be informative, but they often lack important disease-specific signals compared with BAL and airway tissue transcriptomics, underscoring the need for alternative minimally invasive molecular biomarkers for asthma [1]. Severe EA (sEA) is predominantly neutrophilic, whereas mild-to-moderate EA (mEA) may exhibit neutrophilic, mastocytic, eosinophilic or mixed BAL phenotypes. There is also evidence for the occurrence of paucigranulocytic asthma in both humans and horses [2], [3]. Airway remodeling processes involving epithelial barrier dysfunction, smooth muscle hyperplasia, and subepithelial fibrosis have been shown to contribute to disease pathogenesis and are a known feature of sEA, but they have also been implicated in mEA [4], [5], [6].

The heterogeneous nature of EA, together with the invasiveness of current diagnostic approaches, poses significant challenges for disease monitoring and therapeutic intervention. Effective biomarkers are critically important for EA management as they would enable early disease detection before irreversible airway damage occurs, facilitate non-invasive monitoring of disease progression and treatment responses, allow for phenotype-specific therapeutic approaches, and provide insights into local and systemic inflammatory processes. Furthermore, accessible biomarkers could enhance veterinary practice by reducing the need for repeated invasive procedures while improving diagnostic accuracy and enabling personalized treatment strategies for affected horses.

Plasma cfRNA has recently emerged as a promising biomarker platform in oncology, prenatal medicine, and systemic inflammatory research [7], [8]. This circulating transcriptome comprises RNA molecules released through cellular apoptosis, necrosis, and active secretion within extracellular vesicles, reflecting dynamic tissue-specific gene expression profiles [9]. The inherent stability of cfRNA in circulation, conferred by vesicular encapsulation and ribonucleoprotein complex formation, enables its detection and analysis from routine blood samples [10]. The unique properties of cfRNA allow for noninvasive monitoring of tissue-specific molecular signatures, real-time assessment of disease progression, and comprehensive profiling of cellular processes across organ systems [11], [12]. This molecular information has the potential to yield valuable insights into tissue turnover, intercellular communication, and pathophysiological processes that remain undetectable through conventional circulating blood cell analyses [13].

In this study we performed the first characterization of plasma cfRNA in horses, and compared cfRNA with partially matched BAL and whole blood (WB) transcriptomes. The aims were to (i) define the cellular and tissue contributions to the cfRNA pool in both healthy horses and EA, (ii) identify differentially expressed transcripts and pathways, and (iii) characterize shared and distinct gene signatures of cfRNA with BAL and WB to identify unique cfRNA-derived molecular insights into EA.

## Results

### EA samples collection and multiple biological compartments study design

We collected samples from three biological compartments across a clinically recruited cohort and healthy controls to comprehensively characterize the molecular landscape of EA (**Figure 1A**). Plasma cfRNA was profiled in EA cases (n = 16) and healthy controls (n = 12). cfRNA-stabilizing blood collection was used to minimize ex vivo release of platelet- and leukocyte-derived RNA, in combination with a total RNA-seq workflow optimized for low input and incorporating unique molecular identifiers (UMIs) for noise reduction. EA cases included diverse BAL cytology phenotypes, including mastocytic (n = 11, mEA), paucigranulocytic (n = 1, mEA) and neutrophilic (n = 4 sEA) (**Figure 1B, Supplementary Table 1**). For comparative analysis, we sequenced BAL RNA (n = 12: 7 EA cases, 5 controls) and whole blood (WB) RNA (n = 11: 6 EA cases, 5 controls) from partially matched individuals using the same library preparation protocol. (**Figure 1A-B, Supplementary Table 1**). These additional samples similarly represented mixed mastocytic and neutrophilic phenotypes (**Figure 1B**). We obtained full three-compartment sampling from 9 horses, with additional horses contributing two-compartment pairs. (**Figure 1C**). The cfRNA sequencing analysis detected an average of 23,220 genes per sample at ≥1 counts per million (CPM) and 12,497 genes at ≥5 CPM, out of a total of 38,666 genes. The corresponding numbers for BAL/WB were 18024/18613 genes (≥1 counts) and 13264/13544 genes (≥5 counts).

**Figure 1.**
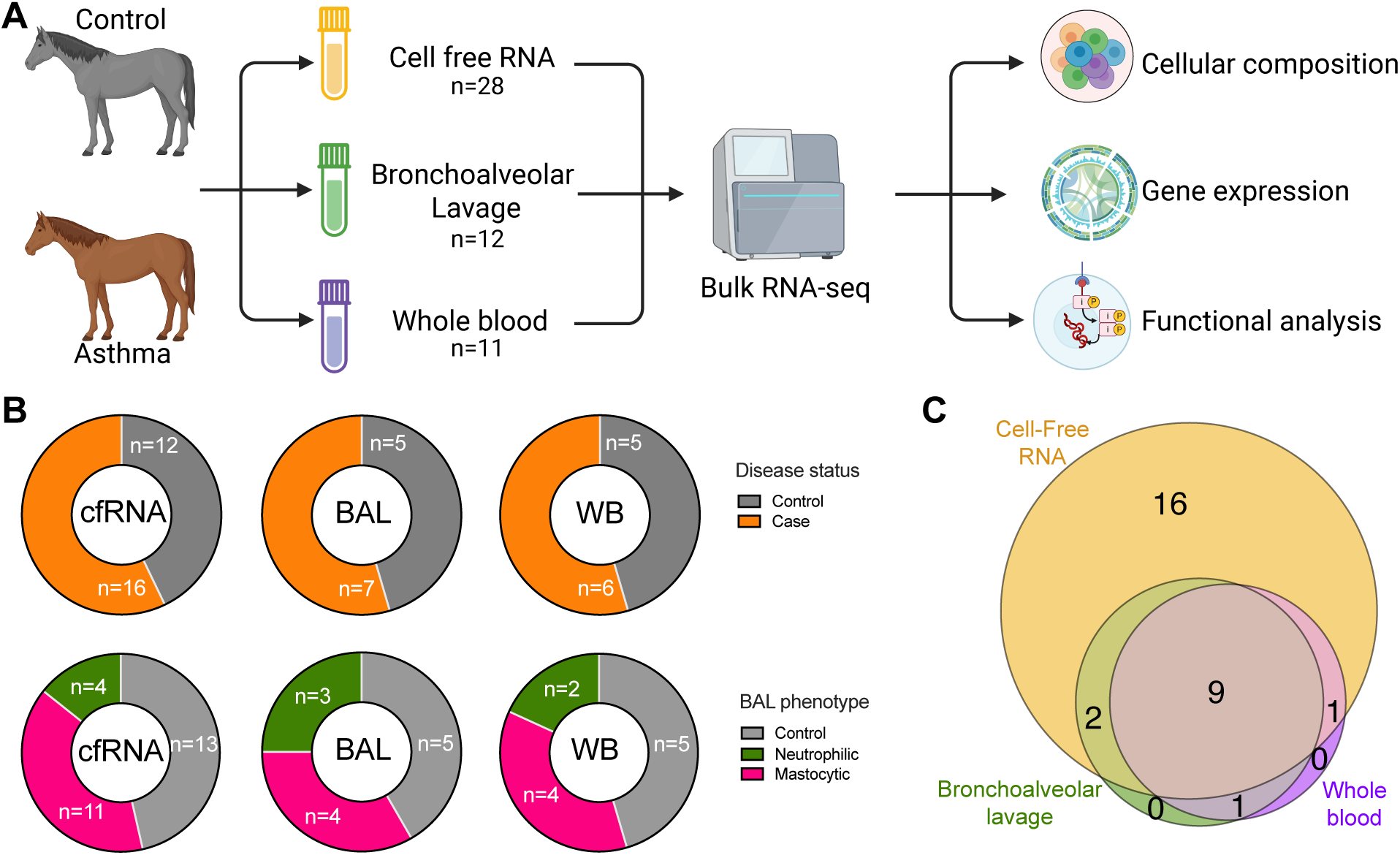
Sample distribution and overlap across biological compartments in equine asthma transcriptomics study. **A) Study design**. Schematic overview of the multi-compartment sampling strategy. **B) Sample size distribution by experimental groups**. Upper panel: Distribution of samples between control (gray) and asthmatic case (orange) groups across compartments. Lower panel: Distribution by BAL cytological phenotype within each compartment, stratified by control (gray), mastocytic (pink), and neutrophilic (green) inflammatory patterns. **C) Sample overlap**. Venn diagram illustrating the intersection of 29 samples collected across the three biological compartments.

### Characterization of the plasma cell-free RNA composition in horses with asthma and controls

To infer tissue of origin, equine genes were mapped to human orthologues and analyzed through deconvolution using the established human pan-tissue reference from Tabula Sapiens (TSP) and the immune cell–specific LM22 reference set [14], [15], [16], [17]. Despite reliance on cross-species references, conservation of tissue markers across mammals supports this approach [18]. Deconvolution analysis revealed that equine cfRNA originated from diverse cellular sources, with immune cells forming the predominant component (B cells, T cells, myeloid and dendritic cells, NK/cytotoxic cells, platelets, neutrophils, ILCs, and a small mast-cell fraction). Signals were also detected from epithelial and stromal tissues, including intestinal epithelium (tuft and goblet cells), endothelial and basal cells, muscle, and peripheral nervous system. Importantly, cfRNA potentially derived from lung ciliated and goblet cells was detectable, indicating respiratory tract contribution to the cfRNA pool (**Figure 2A**).

**Figure 2.**
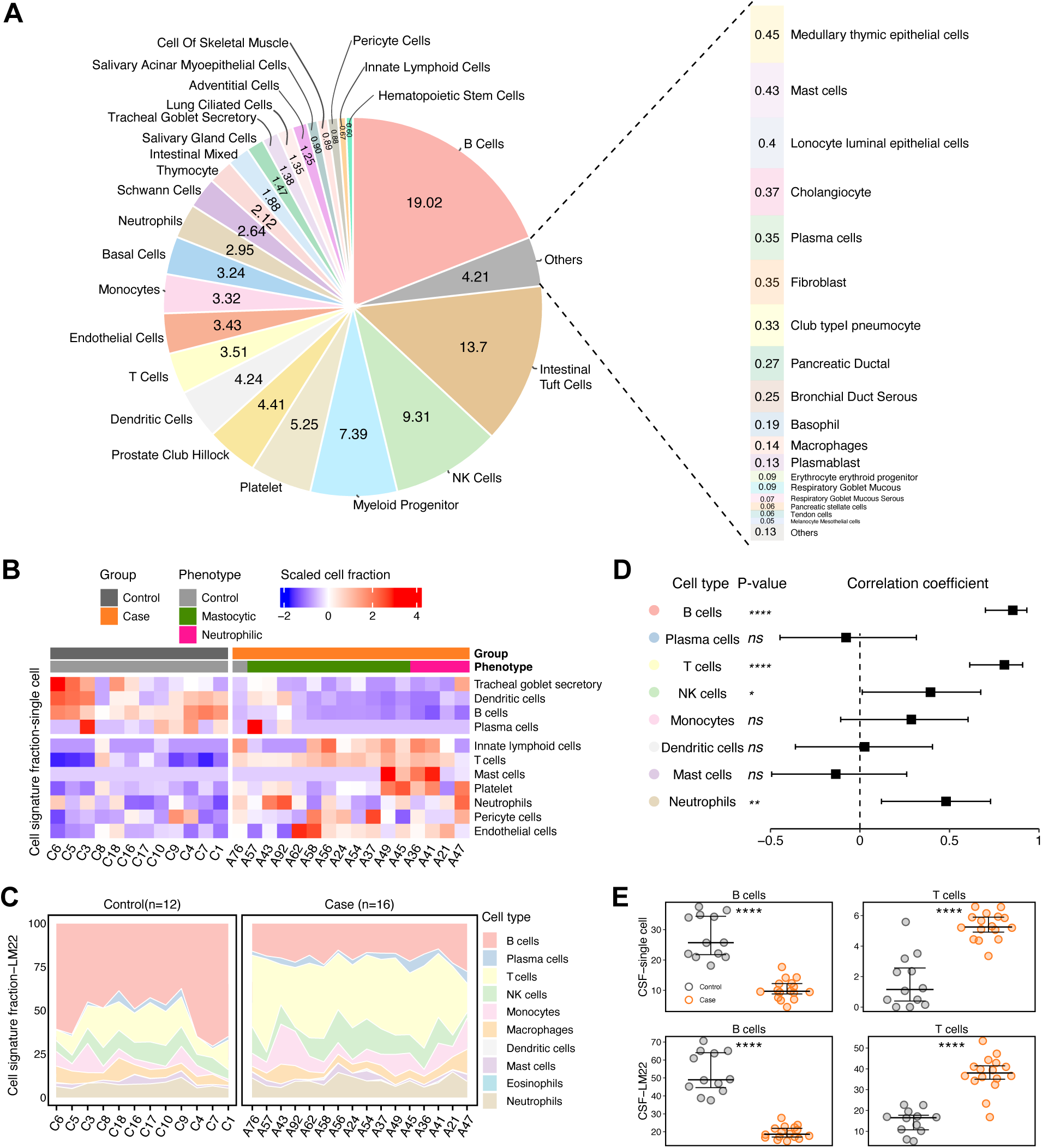
Cell-free RNA deconvolution reveals differential cellular contributions in equine asthma. **A) Comprehensive tissue-of-origin profiling.** Cross-species cell signature deconvolution based on the Tabula Sapiens (TSP) pan-tissue single cell reference data set. **B) Disease-associated alterations in cell-type signatures.** Hierarchical clustering heatmap displaying Z-score scaled TSP cell-type signatures across equine asthma cases and controls (p < 0.05). **C) Leukocyte deconvolution using LM22 reference.** Stacked area plots illustrating the proportional distribution of 10 major immune cell populations estimated using the LM22 leukocyte signature matrix. Sub cell-types were collapsed into their major cell type (B-cells, T cells, etc) **D) Cross-reference validation of deconvolution methods.** Forest plot displaying correlation coefficients between cell fraction estimates derived from TSP and LM22 reference matrices for eight overlapping cell types. Error bars represent 95% confidence intervals. Statistical significance: ****p < 0.0001, **p < 0.01; ns (not significant, p > 0.05). **E) Validation of key immune signature alterations.** Dot plots comparing B cell and T cell fractions between control and case groups using two independent deconvolution approaches: TSP reference (top) and LM22 reference (bottom). Each dot represents an individual sample, with thick horizontal lines indicating group medians (****p < 0.0001).

Comparative analysis of EA cases and healthy controls revealed significant alterations in cellular contributions to the cfRNA landscape. EA samples showed reduced proportions of B-cell and dendritic cell signatures, alongside elevated fractions of T-cell, endothelial cell, and ILC signatures (**Figure 2B, Supplementary Table 2**). The most prominent and consistent findings were reduced B-cell signatures (TSP: p < 10⁻⁴; LM22: p < 10⁻⁴) and increased T-cell signatures (TSP: p < 10⁻⁴; LM22: p < 10⁻⁴) in EA samples (**Figure 2B, C).** Consistent with these observations, strong correlations between the two reference datasets were noted for B cells (r = 0.858, p < 10⁻⁴) and T cells (r = 0.949, p < 10⁻⁴), supporting the robustness of our analytical approach (**Figure 2D**). Furthermore, this immune cell redistribution pattern was preserved across all included EA phenotypes (**Figure 2E**). Leukocyte-focused (LM22) deconvolution additionally revealed increased relative contributions from CD4⁺ naïve and memory T cells, accompanied by concurrent reductions in T follicular helper (Tfh) and memory B-cell signatures (**Supplementary Figure 1A**).

Together, these findings show that circulating cfRNA profiles in horses with EA reflect consistent immune cell redistribution, characterized by reduced B-cell and increased T-cell signatures across phenotypes.

### Differential cfRNA expression in equine asthma revealed upregulation of a key asthma alarmin, remodeling mediators, and immune cell transcripts

To further characterize cfRNA alterations associated with EA, we performed additional transcriptome-wide analyses. Principal component analysis (PCA) indicated separation between EA and control cfRNA profiles (**Figure 3A**). Differential expression analysis identified asthma-associated alarmin *IL33* and *IGHE* among the most significantly elevated transcripts in the EA cohort (log₂FC > 2.5, p < 10⁻⁴) [19]. Upregulation of both these genes is consistent with type-2–skewed inflammation characteristic of allergic airway disease (although cfRNA data and these markers alone are not definitive). Additional upregulations were observed in extracellular matrix (ECM) and remodeling mediators (*MMP1, MMP8, SERPINE1, ADAM9, HPSE, TGFB2, TIMP4*), kinase signaling (*AKT3*), cytoskeletal regulators *(KRT16*) and platelet-associated factors *(APP*) [20], [21], [22], [23], [24], [25], [26], [27], [28]. Increased immune-related transcripts included T-cell activation markers (*CD3D, CD28, CD4, CD226, CD247, LAT, ITK, CD69, IL7R*), NK/cytotoxic molecules (*KLRF1, NCR1, GZMK, GZMH*), and chemokine receptors (*CCR4, CXCR4*) (**Figure 3B**). A list of up- and downregulated genes in cfRNA is provided in **Supplementary Data 1** as well as a compilation of functional themes **(Supplementary Table 3).**

**Figure 3.**
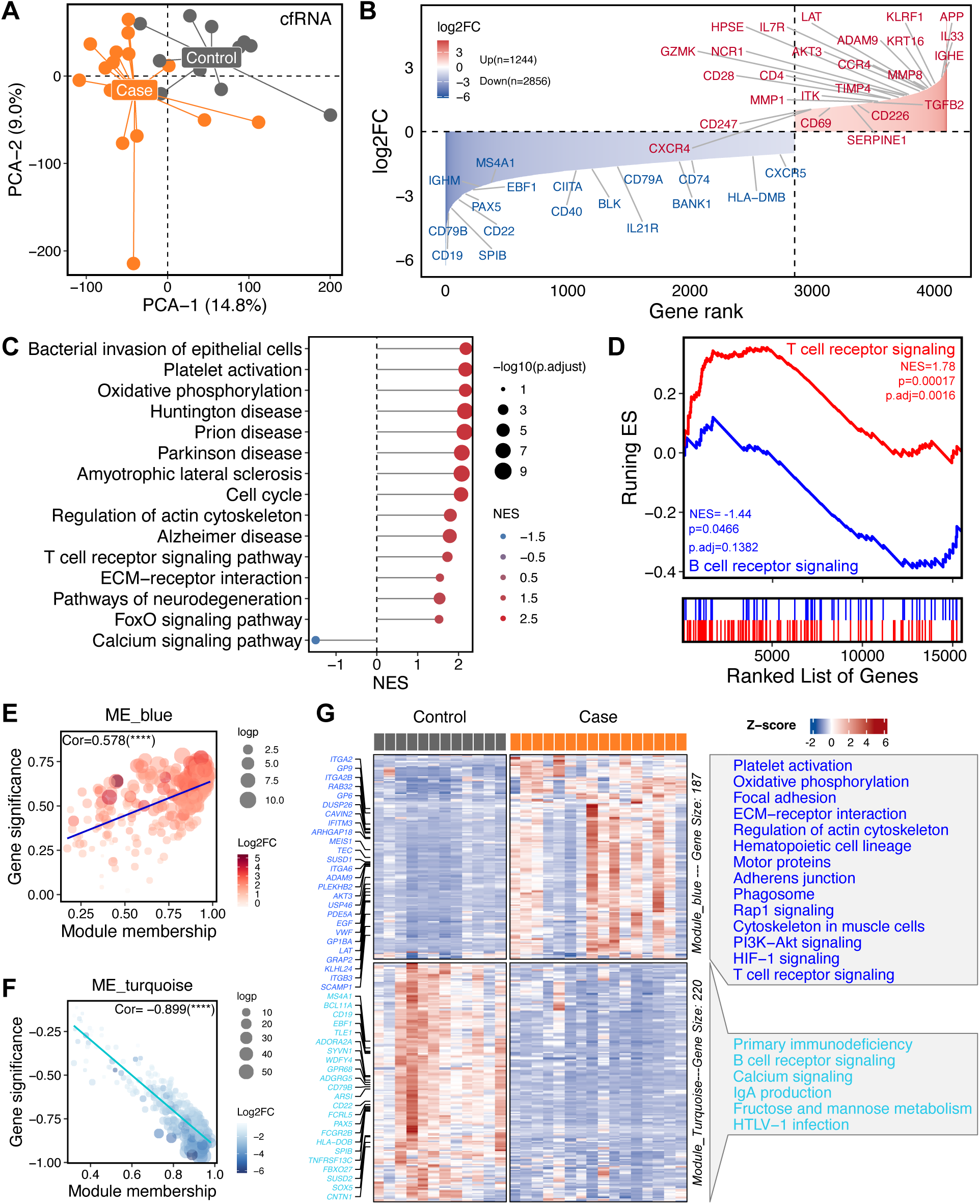
Cell-free RNA transcriptomic profiling reveals disease-specific molecular signatures in equine asthma. **A) Principal component analysis reveals distinct transcriptomic profiles**. **B) Differential gene expression landscape.** 1,244 upregulated genes (red, log₂FC > 1, adjusted p < 0.05) and 2,856 downregulated genes (blue, log₂FC < -1, adjusted p < 0.05. **C) Pathway enrichment analysis.** Lollipop plot showing GSEA results for significantly altered biological pathways (KEGG) between asthmatic cases and controls. **D**) **GSEA plot of the T cell receptor signaling and B cell receptor signaling pathway. E & F**) **Weighted gene co-expression network analysis (WGCNA).** The two most significant co-expressed modules: *module_blue* (upregulated) & *module_turquoise* (downregulated). Correlation plots between gene significance and *module_blue* membership gene significance and *module_turquoise* membership**. G**) Heatmap of genes in *module_blue* and *module_turquoise*. Potential asthma-related genes are highlighted on the left, with corresponding over-representation analysis results shown on the right.

In contrast, B-cell developmental and functional programs showed reduced representation in EA cfRNA samples. This included canonical B-cell markers (*MS4A1, CD19, CD79A/B, CD22*), lineage regulators (*PAX5, EBF1, SPIB*), proximal signaling components (*BANK1, BLK),* selected Fc receptor family members, and antigen-presentation/co-stimulatory molecules (*CIITA, DMB, CD74, CD40*)[29], [30]. In addition, Tfh-associated transcripts (*CXCR5, IL21R*) were lower, consistent with a diminished Tfh-related signal [31]. Concordantly, multiple immunoglobulin locus transcripts (*IGHM, IGHV, IGLC3*) were reduced (**Figure 3B).**

Collectively, these patterns confirm the results from the deconvolution analysis and suggest a relative attenuation of B-cell/Tfh transcriptional signals in plasma cfRNA in the context of type-2–skewed inflammation, ECM remodeling, and heightened T-cell activity-This is while also acknowledging that cfRNA reflects circulating transcript release rather than direct cellular function.

Pathway enrichment of cfRNA DEGs revealed enrichment of transcripts consistent with processes relevant to asthma pathophysiology, including oxidative stress responses, epithelial defense, tissue remodeling, platelet activation, and immune cell signaling (**Figure 3C)**. Interestingly, pathways annotated as neurodegenerative disease (Prion, Huntington, ALS, Parkinson, Alzheimer, pathway of neurodegeneration) were significantly enriched (NES > 1, p < 0.05). These signals likely reflect shared mitochondrial and oxidative stress rather than neuron-specific degenerative pathology. Key alterations included a pronounced up-regulation of oxidative phosphorylation, enrichment of bacterial-invasion–of– epithelial-cells pathways, and regulation of the actin cytoskeleton—consistent with epithelial-barrier activation and cytoskeletal remodeling. Additional enriched categories encompassed cell cycle regulation, ECM-receptor interactions, FoxO signaling, and calcium signaling pathways. Gene set enrichment analysis (GSEA) supported the immune cell redistribution observed in the deconvolution results, showing significant enrichment of T-cell receptor signaling pathways (NES = 1.78, p = 0.0016) and a trend toward depletion of B-cell receptor signaling networks, although this did not reach statistical significance after multiple testing correction (NES = –1.44, p = 0.1382) **(Figure 3D**).

Furthermore, weighted gene co-expression network analysis (WGCNA) identified two distinct modules (MEs) among six modules that associated with EA disease status (**Supplementary Figure 2A, Supplementary Table 4**). The most significant co-upregulated module (ME_blue, n = 187 genes) was positively correlated with EA cases (r = 0.578, p < 10⁻⁴) and included genes related to platelet activation, oxidative phosphorylation, focal adhesion, and ECM–receptor interactions (**Supplementary Figure 2B, Supplementary Table 4**). This module was further enriched for pathways regulating actin cytoskeleton dynamics, hematopoietic cell lineage specification, and adherens junction formation, as well as signaling cascades such as Rap1, PI3K–Akt, HIF-1, and T-cell receptor signaling (**Figure 3E, G**). Conversely, the co-downregulated module (ME_turquoise, n = 220 genes) showed a strong negative correlation with disease status (r = –0.899, p < 10⁻⁴) and was enriched for genes involved in primary immunodeficiency, B-cell receptor signaling, calcium signaling, and immunoglobulin A (IgA) production pathways (**Figure 3F, G, Supplementary Figure 2C)**.

Principal component analysis (PCA) did not reveal significant group-level separation of cfRNA transcriptomes among the different EA/BAL phenotypes included in this study (**Supplementary Figure 3A**). Given the pilot nature of this study and the need for increased statistical power, we limited our main analysis to the joint cohort. Nevertheless, differential gene expression (DGE) and gene set enrichment analysis (GSEA) for mastocytic (n = 11) and neutrophilic (n = 4) cases versus controls are provided in **Supplementary Figure 3** and **Supplementary Data 1.**

### Multi-tissue transcriptomic analysis reveals compartment-specific molecular signatures and alterations in equine asthma

Next, we observed expression patterns in two additional biological compartments in a subset of horses with EA vs controls. BAL transcriptomes from EA horses (n = 7) showed induction of multiple inflammatory and some tissue remodeling genes. Upregulated genes included pro-inflammatory mediators *(OSM, IL20RA, CCR10*), feedback regulators (*CRISPLD2, DUSP1*), immediate-early transcriptional regulators (*EGR1, FOS/FOSB, MS4A4A/7*), and mast-cell/epithelial interaction markers (*MRGPRX2, AREG*) [32], [33],[34], [35], [36], [37]. Structural/remodeling-associated genes (*TGFB3, MMP8, ACTA2*) were increased, as were innate antimicrobial transcripts such as *LYZ*, and neuro-immune signaling transcripts (*NGF, RGS2, RGS8, GRIK4, GRIN1, GABRR1*) (**Figure 4A–B**). KEGG analysis identified enrichment of (i) Neuroactive ligand– receptor interaction (NES = 1.32, FDR-adjusted p = 0.0197), consistent with increased ligand–receptor signaling spanning neuro-immune and intercellular communication pathways, and (ii) Adrenergic signaling in cardiomyocytes (NES = 1.37, FDR-adjusted p = 0.0213), possibly reflecting β-adrenergic signaling relevant to airway smooth muscle tone (despite the cardiomyocyte-centric KEGG label) (**Figure 4C**). [38][39]. Separate DGE analysis of neutrophilic (n = 3) and mastocytic (n = 4) cases were not sufficiently powered due to the low numbers, but are nevertheless shown in **Supplementary Figure 4**.

**Figure 4.**
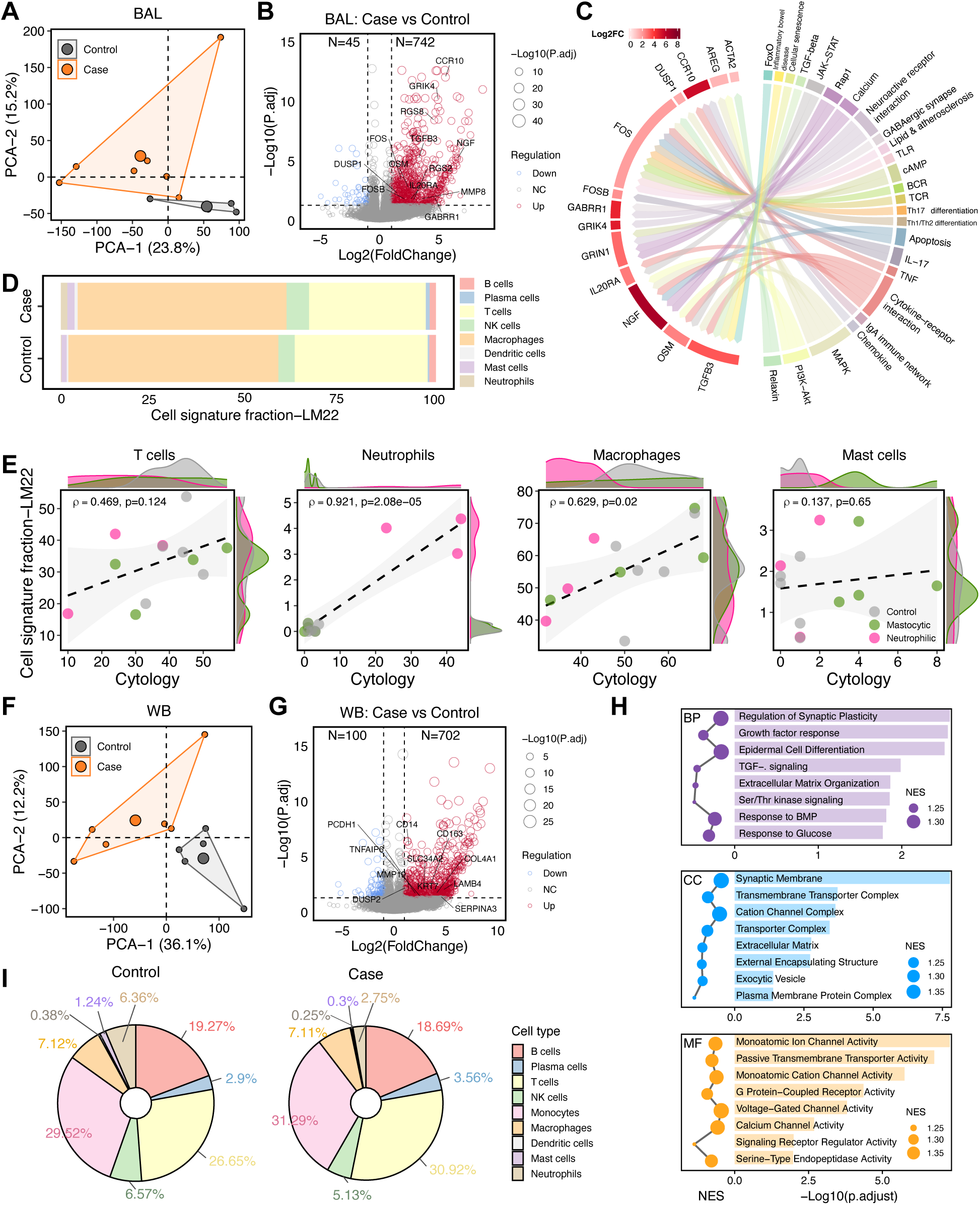
Transcriptomic landscapes of BAL and whole blood compartments. A**) PCA analysis of the BAL transcriptome landscape. B) BAL differential gene expression profile.** Volcano plot illustrating 745 upregulated vs 45 downregulated genes in equine asthma cases. **C) Pathway activation network in BAL.** Chord diagram illustrating significantly expressed genes and associated KEGG pathways in asthmatic versus control BAL samples. Ribbons connect pathways (outer arc) to their constituent differentially expressed genes (inner arc), with color intensity representing log₂ fold change magnitude. **D) Comparative BAL cell composition in EA vs controls.** Stacked bar chart displaying the average relative proportions of eight major immune cell types in BAL samples deconvoluted using the LM22 reference. **E) Validation of computational deconvolution fractions against cytology fractions.** Scatter plots correlating computationally estimated cell fractions (LM22, y-axis) with presumed ‘ground truth’ cytological measurements (x-axis) for four cell types. Each point represents an individual BAL sample, colored by cytological phenotype (gray: control, pink: mastocytic, green: neutrophilic). **F) PCA analysis of the whole blood transcriptome landscape. G) Whole blood differential expression landscape.** Volcano plot illustrating 702 upregulated vs 100 downregulated genes in equine asthma cases. **H) Gene Ontology pathway enrichment analysis.** Dot plots displaying significantly enriched biological processes (BP), cellular components (CC), and molecular functions (MF) in whole blood samples from asthmatic cases versus controls. Dot size represents the normalized enrichment score (NES), while the length of bar indicates statistical significance (-log₁₀ adjusted P-value). **I) Comparative whole blood cell composition in EA vs controls.** Pie charts illustrating the relative proportions of major immune cell populations in WB samples from control and asthmatic case groups.

Cellular deconvolution analysis, using either the LM22 reference matrix or an equine BAL scRNA-seq derived reference, revealed no significant differences in overall cell type proportions between EA cases and controls (**Figure 4D, Supplementary Figure 5)** [40]. Consistent with previous observations from scRNA-seq experiments, the proportional estimates of neutrophils and mast cells in BAL deconvoluted transcriptomic data did not exactly match cytology measurements. However, neutrophil fractions showed strong correlation with cytological assessments (ρ = 0.921, p = 2.08 × 10⁻⁵) whereas mast cell fractions did not (**Figure 4E**).

Whole blood EA transcriptomes (n = 6) also revealed systemic molecular signatures consistent with inflammatory activation and signals associated with extracellular-matrix remodeling. Upregulated genes included innate immune and monocyte-macrophage markers (*CD14, CD163, TNFAIP6, DUSP2, SERPINA3*) alongside matrix-remodeling enzymes (MMP19) indicating systemic inflammatory responses. Notably, airway- and barrier-associated transcripts (*PCDH1, KRT7, SLC34A2, LAMB4, COL4A1*) were elevated, suggesting epithelial injury and subsequent release of airway-derived RNA into circulation (**Figure 4G**) [41], [42], [43], [44]. Gene Ontology (GO) enrichment showed epidermal cell differentiation (potentially reflecting epithelial repair signals), TGF-β/BMP signaling, and ECM organization (**Figure 4H**). By contrast, neuroactive ligand–receptor interaction was the only significantly enriched KEGG pathway in whole blood, mirroring the BAL signal and implicating shared ligand–receptor–mediated (neuro-immune) signatures across compartments (**Supplementary Figure 6A**).

Finally, cellular deconvolution analysis of whole blood samples using LM22 and an equine single cell PBMC reference matrices [60] showed no significant differences in immune cell composition between EA cases and controls, consistent with BAL findings (**Figure 4I, Supplementary Figure 6B-C**).

### Comparative transcriptomic profiling across biological compartments identifies tissue-specific and overlapping molecular signatures in EA

We next compared transcriptional profiles across the three biological compartments to assess compartment-specific and overlapping disease signatures in EA. PCA revealed distinct transcriptional landscapes (**Figure 5A**). Cross-compartment correlation analysis of differential expression statistics showed weak associations between compartments (**Figure 5B**). cfRNA exhibited minimal negative correlation with both BAL (r = -0.06, p < 0.0001) and whole blood (r = -0.11, p < 0.0001), while BAL and whole blood showed modest positive correlation (r = 0.32, p < 0.0001). Consistent with the modest positive correlation observed between BAL and WB at the global level, 374 genes (369 upregulated and 5 downregulated) were significantly differentially expressed in both compartments. These overlapping DEGs exhibited highly concordant changes in both direction and magnitude (ρ = 0.82, p < 0.0001), indicating coordinated transcriptional responses between the airway and peripheral blood (**Figure 5C**). However, these genes did not show similar expression patterns in cfRNA. Among the 369 genes upregulated in both BAL and WB, only 21 were also significantly upregulated in cfRNA, whereas 95 were significantly downregulated and 253 were not significantly changed. Of the five genes downregulated in both BAL and WB, one was significantly upregulated, one significantly downregulated, and three remained unchanged in cfRNA. While cfRNA exhibited limited gene sharing with either compartment, reinforcing its distinct molecular profile, a small number of transcripts were consistently upregulated across all three compartments, including structural proteins (SMTN, ACTG2), tissue remodeling factors (THSD4), and the stress-response regulator FKBP5 (**Figure 5D**). Notably, three small nucleolar RNAs (SNORD45B, SNORD12, SNORD58A) were consistently more abundant across cfRNA, BAL, and WB, suggesting systemic dysregulation of RNA processing machinery (**Figure 5D).** More broadly, large numbers of snoRNAs were differentially expressed in all three compartments (cfRNA: 71, BAL: 77, WB: 37), indicating a widespread perturbation of noncoding RNA expression and RNA processing pathways in EA (**Supplementary Data 1**).

**Figure 5.**
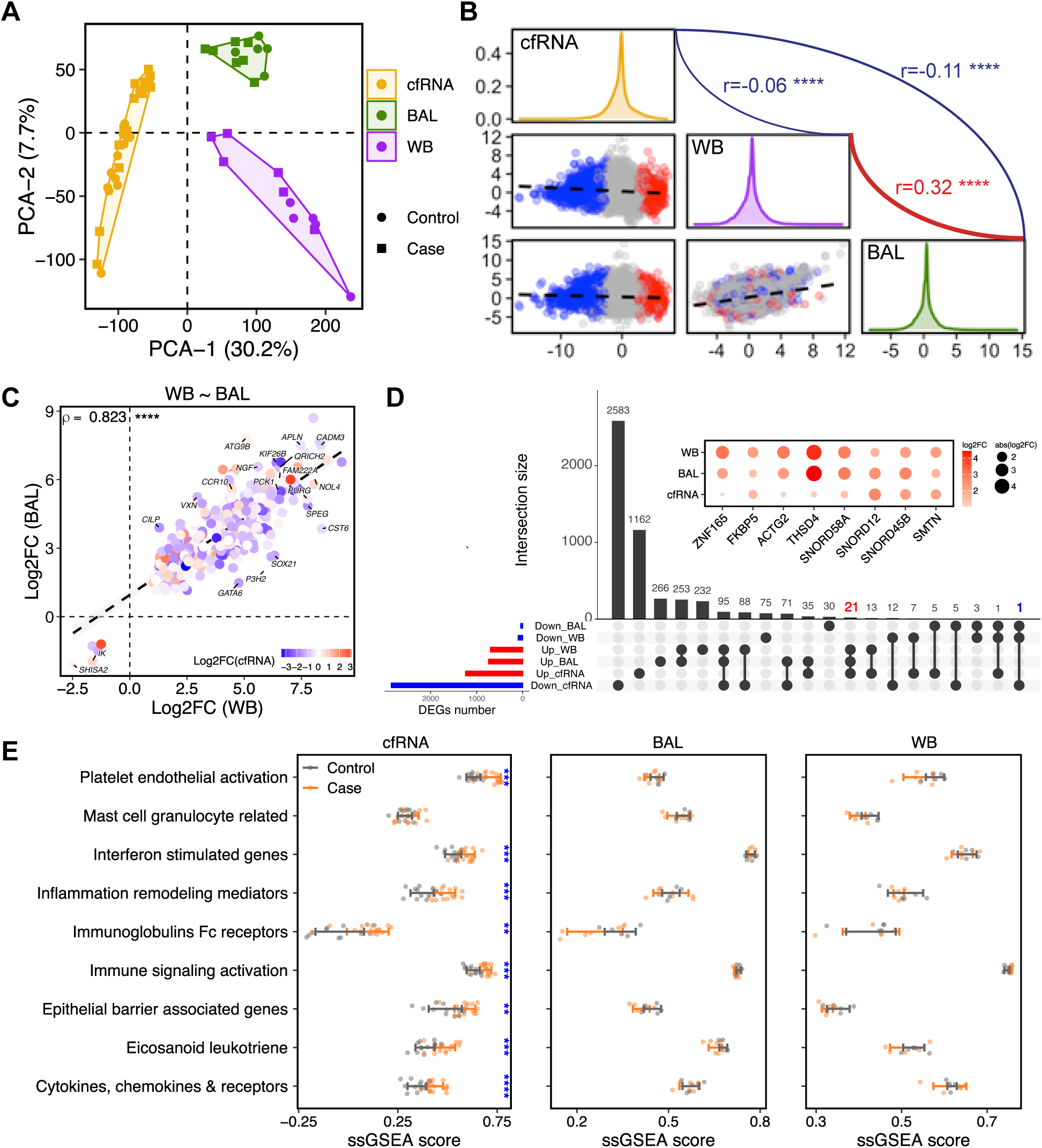
Cross-compartment analysis of molecular signatures in cfRNA, BAL and whole blood. **A)** PCA of TPM value demonstrates clear separation across the three compartments along PC1 (30.2%) and PC2 (9.0%). **B) Inter-compartment correlation of differential expression patterns.** Pairwise scatter plots display DESeq2 stat value between compartments. Each point represents a gene, colored by its log2FC in cfRNA. Density plots show the *stat* value distributions within each compartment. Correlation coefficients between compartments are indicated in the upper-right corners**. C) Correlation of log₂FC for shared DEGs between WB and BAL**. Scatter plot showing genes that exhibited concordant up-or downregulation in both WB and BAL samples when comparing disease and control groups. The color of each point indicates the corresponding gene’s log₂ fold change in cfRNA. **D) Comprehensive intersection analysis of differential expression and cross-compartment biomarker expression profiles.** UpSet plot quantifying the overlap patterns of DEGs across all three sample types and regulation directions. Dot plot displaying log₂FC for nine genes consistently differentially expressed across all three biological compartments. **E) Single-sample Gene Set Enrichment Analysis (ssGSEA) scores for nine asthma-related functional modules across the three compartments.** All modules except the Mast Cell Granulocyte Related module (ssGSEA score: mean (Case) = 0.30, mean (Control) = 0.28; Wilcox test: p = 0.5, p adj = 0.5) were upregulated in EA cfRNA.

To further assess compartment-specific detection of asthma-relevant biology, we analyzed nine predefined gene modules representing processes associated with asthma, selected based on the pathways found upregulated in cfRNA [45], [46], [47], [48], [49]. Modules were curated from established asthma biology and cfRNA-enriched pathways to capture core processes relevant to EA. The module genes are provided in **Supplementary Table 5**. In this partially matched cohort, cfRNA showed broader pathway-level transcript perturbations than BAL or whole blood. Using gene-level differential expression and ssGSEA (p < 0.05), cfRNA from EA cases showed significant upregulation of multiple modules, including platelet– endothelial activation, interferon-stimulated genes, inflammatory/remodeling mediators, innate/adaptive immune signaling, epithelial barrier–associated genes, eicosanoid/leukotriene pathways, and cytokine– chemokine receptor networks (**Figure 5E**, **Supplementary Table 5**) [50]. By contrast, BAL and WB did not show significant enrichment of these modules between cases and controls (**Figure 5E**). These findings indicate that each compartment captures fundamentally different aspects of EA pathophysiology, with cfRNA uniquely capturing molecular perturbations that were essentially undetectable in conventional sampling approaches. However, because module composition partly reflects cfRNA results, cross-compartment sensitivity comparisons should be considered exploratory and may favor cfRNA.

## Discussion

cfRNA provides a dynamic readout of tissue turnover and extracellular signaling rather than a snapshot of circulating cell counts, because it reflects transcripts released by dying cells or exported in extracellular vesicles. This can capture active processes, cell loss and replacement and intercellular communication, that may not be apparent from conventional blood or BAL measurements. However, because cfRNA reflects RNA release, packaging, and clearance, cfRNA abundance should be interpreted as enrichment of circulating transcript signals consistent with biological processes rather than as direct evidence of transcriptional activation in source tissues.

Although circulating lymphocytes contribute to cfRNA, they represent only a small share of the total lymphocyte pool (∼2% in humans). Most lymphocytes reside in peripheral barrier tissues; lung, gastrointestinal tract, and skin each harbor ∼3–4%, so a substantial fraction of lymphocyte-derived cfRNA likely originates from tissue-resident populations [51]. Consistent with human plasma cfRNA deconvolution work, this study found lymphocyte-derived signals alongside transcripts specific to intestinal organs, skeletal muscle, and lung tissue, underscoring the promise of plasma cfRNA analysis for equine conditions beyond asthma (e.g., colic and ulcers). Because cfRNA reflects RNA release, packaging, and clearance, cfRNA abundance should be interpreted as enrichment of circulating transcript signals consistent with biological processes rather than as direct evidence of transcriptional activation in source tissues.

cfRNA transcriptomics revealed disease-associated processes in equine asthma (EA) that were not evident or clearly enriched in bulk WB or BAL transcriptomes, including altered lymphocyte balance, epithelial alarmin signals, and structural remodeling. The most prominent immune feature was a reduction in B-cell signatures, accompanied by increased CD4⁺ T-cell and innate lymphoid cell (ILC) signatures and elevated *IL33* and I*GHE* transcripts. The pattern is consistent with an IL-33–associated, CD4⁺ T-cell/ILC-skewed type-2 milieu detectable by cfRNA in this cohort. IL-33 protein is abundant in the nuclei of endothelial, epithelial, and stromal cells and is released upon tissue damage. Increased detection of IL-33 mRNA in plasma-derived RNA could represent a surrogate marker of epithelial activation or tissue injury in equine asthma. The role of IL-33 in allergic asthma is well established, however, it has also been implicated in non-allergic inflammation and in COPD in humans [52].

The concomitant reduction in B-cell and Tfh-associated transcripts, together with the absence of upregulation in follicular chemokines (CXCL13, CCL19, CCL21) in cfRNA, suggests limited systemic B-cell or follicular activity. Alternatively, this attenuated cfRNA signature may indicate compartmentalization of B-cell responses within inducible tertiary lymphoid structures in the lungs (e.g. iBALT) during inflammation, potentially resulting in reduced cfRNA release into circulation.

This study cohort comprised a mix of mEA and neutrophilic sEA, which may introduce variability in the observed transcriptional profiles. Recent single-cell studies underscore the heterogeneity of EA, suggesting Th17 polarization and B-cell expansion in severe neutrophilic phenotypes [61]. However, type 2 and type 17 programs may co-exist within individuals and vary with disease stage or phenotype [53]. Our findings therefore complement these observations, suggesting that some EA phenotypes are type-2-biased, whereas neutrophilic sEA has previously been found to be predominantly Th17-skewed.

Beyond immune shifts, cfRNA signatures indicated remodeling-related programs, with increased transcripts linked to epithelial barrier activation/repair, matrix turnover, and growth-factor signaling, alongside platelet–endothelial activation and oxidative phosphorylation [54]. Smooth-muscle–associated cytoskeletal transcripts (ACTG2, SMTN) were consistently upregulated across cfRNA, BAL, and whole blood, as were THSD4 (ECM assembly) and FKBP5 (stress response), supporting active tissue remodeling and cellular stress adaptation [55], [56]. These molecular features align with established histopathology in equine and human asthma—epithelial hyperplasia, submucosal thickening, and smooth-muscle remodeling, and position cfRNA as a minimally invasive surrogate for monitoring airway structural change.

This study is not without limitations. First, the cohort size was modest, which limits statistical power and prevented phenotype-stratified analyses. Second, cfRNA, BAL, and whole-blood samples were partially matched across individuals, which can dampen apparent cross-compartment concordance. Third, EA cases reflected mixed phenotypes, potentially obscuring phenotype-specific signals. Fourth, cell/tissue deconvolution relied mainly on cross-species references, introducing possible annotation bias. Finally, control horses were a homogeneous group of Standardbred trotters, whereas clinical cases spanned multiple breeds and ages. Using a uniform control group reduces within-group variance and can increase power, but it also limits generalizability and may introduce breed confounding.

## Conclusion

Despite these limitations, our data provide an initial framework for understanding the circulating transcriptome in equine asthma and healthy horses. Plasma cfRNA captured disease-associated immune shifts together with remodeling-related programs. These signatures align with established aspects of asthma pathobiology and were only partially reflected in BAL or whole blood, supporting cfRNA as a minimally invasive, complementary readout of airway and systemic processes. Taken together, the findings nominate cfRNA modules and transcripts as candidate biomarkers for phenotype characterization and for monitoring airway injury/remodeling and inflammatory tone. Future work should validate these signals in larger, phenotype-resolved and longitudinal cohorts; integrate cfRNA with clinical and physiological measures; and employ equine single-cell references for cell-type deconvolution.

## Methods

### Horses and sampling procedures

Healthy control horses were sampled in spring (April) at a national education and harness racing facility (Wången). Control horses passed a detailed clinical examination and endoscopic examination, had no history or signs of any respiratory disease and BAL cytology did not reveal any pathological findings. Case horses diagnosed with equine asthma were sampled during consultancy at an equine referral veterinary hospital (University Animal Hospital, SLU, Uppsala, Sweden). Equine asthma diagnosis was based on BAL cytology (**Supplementary Table 1**), history of symptoms and clinical examination. Bronchoalveolar lavage and BAL cytology was performed as previously described [59]. Whole blood and plasma were collected on the same sampling occasion as BAL from the jugular vein using Tempus Blood RNA tubes (Thermo Fisher) and Streck RNA Complete BCT tubes (Streck). The Tempus tubes contain reagents that lyse and stabilize blood cells immediately upon collection, preserving intracellular RNA. Filled tubes were stored at -80°C until processing. Streck RNA Complete BCT tubes contain stabilizing agents that preserve the integrity of nucleated blood cells and prevent their lysis, as well as maintaining the quality of cell-free RNA in plasma. Plasma was extracted from the Streck tubes according to the manufacturers instruction and stored at -80°C until processing. Ethical permissions (#5.2.18-4819/15, #31-2605/04) was granted by the regional ethical board.

### RNA extraction

Total RNA was extracted from BAL cells using the Qiagen AllPrep Kit. Whole blood (WB) RNA was isolated from Tempus tubes using Norgen’s Total RNA Purification Kit. cfRNA was extracted from 2 ml of plasma using the Quick-cfRNA Serum & Plasma Kit (Zymo Research). RNA integrity was checked on a TapeStation system (Agilent) using RNA Screen Tapes (BAL and WB, RIN > 8) or High Sensitivity RNA Screen Tapes (cfRNA, RIN ≤ 3). cfRNA yields were, as expected, very low (1-5 ng total RNA/ml plasma) and dominated by short molecules. cfRNA yields were generally too low to be reliably quantified by TapeStation or Qubit analysis so the sample volume was reduced by vacuum centrifugation, and the entire concentrate was used for library preparation. All RNA samples, including cfRNA, were DNase treated just prior to library prep using the Heat&Run gDNA removal kit (ArcticZymes).

### RNA-seq library preparation

RNA-seq libraries were prepared using the SMARTer Stranded Total RNA-Seq v3 – Pico Input Mammalian Kit (including UMIs), or the v4 version, renamed SMART-Seq® Total RNA Pico Input with UMIs (ZapR® Mammalian), which according to the manufacturer (Takara Bio) is the same kit as v3 but exhibits improved rRNA removal efficiency. The fragmentation step was omitted for the cfRNA samples. A total of 10 ng of RNA input was used for BAL and WB samples (maximum input specified for this kit). For cfRNA, the entire amount extracted from 2 ml of plasma was used as input (estimated at ∼1–5 ng based on TapeStation profiles). Libraries were sequenced as paired-end 150 bp (PE150) reads on a NovaSeqX instrument. The reason for switching to the new version of the kit (v4) was that the removal of equine ribosomal RNA proved rather inefficient with the v3 kit. However, we did not observe significant improvement in rRNA removal efficiency with the v4 kit. Principal component analysis indicated that batch effects across the v3 and v4 library prep kits were insignificant (data not shown).

### RNA-seq read processing

Best practice alignment, UMI deduplication and quantification was performed with the nf-core/RNA-seq pipeline v.3.8.1 (https://github.com/nf-core/rnaseq, using EquCab 3.0 annotation version 0.111. The number of mapped reads (STAR v.7.10a) were on average 16.83 ± 9.1 million reads for the cfRNA samples, 33.9 ± 8.8 for the WB samples and 33.7 ± 16.6s for BAL samples. UMI deduplication was performed with UMI-tools within the nf-core RNA-seq pipeline (v.1.1.2). Ribosomal reads were higher than expected (Subread_featurecounts v2.0.1), considering the library kit is designed for mammalian rRNA removal: average 39.9 % in cfRNA, 32.7 % in WB and 29.9 % in BAL-data. (**Supplementary Data 2**).

### Deconvolution analysis

Cell deconvolution analysis was performed on equine bulk RNA sequencing data to determine relative cellular contributions. Cross-species ortholog mapping was performed using the biomaRt R package (v2.64.0) to establish correspondence between equine and human gene annotations [63]. To ensure high-confidence cross-species comparisons, stringent filtering criteria were applied, retaining only orthologous relationships with a confidence score of 1, which successfully mapped 15,800 equine genes to their human orthologs. Cell type proportion estimation was performed using validated single-cell reference matrices specific to each sample type. cfRNA analysis utilized the established human tsp_v1_basisMatrix (Tabula Sapiens, TSP) and the human immune cell LM22 signature matrix. BAL cell deconvolution was performed using LM22 as well as a internally generated equine respiratory cell dataset (PRJNA914226) For whole blood analysis LM22 and a reference signature from GSE148416 equine PBMC data was employed [57], [58], [59]. Single-cell datasets were preprocessed using Seurat v5.3.0 with Harmony batch correction and annotated through unsupervised clustering combined with differential expression analysis and established cell type markers [60], [61]. Deconvolution was implemented via CIBERSORTx with signature matrix generation followed by cell fraction imputation using S-mode batch correction, with both reference and bulk RNA-seq data normalized to CPM format [62]. Methodological concordance was assessed through Spearman correlation analysis with significance determined at p.adj < 0.05 and correlation coefficients ≥ 0.5 indicating strong agreement.

### Differential Gene Expression and Pathway Analysis

Raw count matrices representing mRNA expression profiles were generated from aligned sequencing reads across all samples. The initial dataset encompassed 38,849 genes. To ensure analytical rigor, ribosomal genes were excluded, retaining 38,666 genes for downstream analysis. Differential gene expression analysis was conducted independently for each experimental dataset using the DESeq2 R package (v.1.48.1)[64]. Genes were classified as significantly differentially expressed based on stringent statistical criteria: Benjamini-Hochberg adjusted p-value < 0.05 and absolute log₂ fold change ≥ 1.0, corresponding to a minimum two-fold expression change.

To elucidate the biological significance of transcriptional changes, functional enrichment analyses were performed using two complementary methodological approaches implemented in the clusterProfiler R package (v.4.16.0): Over-representation analysis (ORA) was applied to significant DEGs and custom gene sets of interest. Gene Set Enrichment Analysis was conducted on the full ranked gene lists ordered by log2 fold change values, with enrichment scores normalized to determine pathway activation or suppression [65]. Both analytical approaches leveraged comprehensive pathway databases including the Kyoto Encyclopedia of Genes and Genomes (KEGG: https://www.genome.jp/kegg/) and Gene Ontology (GO: https://geneontology.org/docs/go-enrichment-analysis). Pathways with adjusted p-value < 0.05 were considered significantly enriched.

### Weighted gene co-expression network analysis

To explore biological relationships among DEGs and reduce dimensionality, Weighted Gene Co-expression Network Analysis (WGCNA) was performed on bulk cfRNA-seq data to identify co-expression modules associated with disease status [66]. Raw count data were filtered to retain genes with counts > 1 in at least three samples, followed by removal of genes with variance below the 25th percentile to eliminate low-information features. The filtered count matrix was then normalized using DESeq2’s variance-stabilizing transformation (VST). To focus on the most biologically informative and variable genes while maintaining computational tractability, genes with variance above the 95th percentile were selected for network construction, yielding approximately 1,000 genes. WGCNA was performed using a soft-thresholding power of 12, determined by scale-free topology criterion, resulting in the identification of six distinct co-expression modules. Module-trait correlation analysis was conducted to assess the relationship between each module and asthma status. The Module_ turquoise exhibited significant negative correlation with asthma, indicating that genes within this module were predominantly downregulated in asthmatic horses relative to controls, while the Module_ blue showed significant positive correlation with asthma, suggesting upregulation of its constituent genes in diseased animals. Genes from these two disease-associated modules were extracted for subsequent expression pattern visualization and KEGG pathway over-representation.

### Comparative transcriptomic profiling

To evaluate transcriptomic concordance and divergence across cfRNA, BAL, and WB compartments, integrative comparative analyses were performed. Principal component analysis (PCA) was conducted using transcript-level expression values (TPM) for all detected genes to visualize global expression patterns and assess sample clustering across compartments. Stat values -DESeq2-derived Wald statistics reflecting both the direction and significance of gene expression changes—were extracted from independent differential expression analyses of each compartment. The distributions of stat values were compared, and pairwise Pearson correlations were calculated to characterize their respective transcriptional response profiles. Additionally, the overlap of DEGs (adjusted p-value < 0.05, |log₂FC| > 1) across the three compartments was calculated to identify shared and compartment-specific transcriptional signatures associated with equine asthma.

### Customized asthma-related functional modules construction

To overcome the broad and generalized annotations of standard KEGG and GO databases, nine customized asthma-related functional modules were constructed to capture biologically relevant pathways implicated in asthma pathogenesis. These modules encompassed: Cytokines, Chemokines & Receptors (n=12 genes); Immunoglobulins & Fc Receptors (n=3); Inflammation & Remodeling Mediators (n=13); Epithelial & Barrier-Associated Genes (n=5); Mast Cell & Granulocyte-Related Genes (n=6); Eicosanoid/Leukotriene Pathway Genes (n=7); Immune Signaling & Activation (n=14); Platelet/Endothelial Activation (n=13); and Interferon-Stimulated Genes (n=11). Module composition was based on established literature and functional roles in asthma immunopathology. Single-sample gene set enrichment analysis (ssGSEA) was performed to quantify pathway-level activity for each module across individual samples, generating sample-specific enrichment scores [67]. Module scores were compared between asthmatic and control horses using Wilcoxon rank-sum tests to identify modules exhibiting differential activation or suppression under disease conditions.

### Statistical analysis

All statistical analyses were conducted using R version 4.5.0 (https://www.r-project.org). Prior to statistical testing, data distributions were assessed for normality using the Shapiro-Wilk test. For data following a normal distribution, Student’s t-test for two-group comparisons were applied; for data not meeting normality assumptions, Wilcoxon rank-sum test (Mann–Whitney U test) were used instead. When multiple comparisons were performed, p-values were adjusted using the Benjamini-Hochberg procedure to control the false discovery rate. Adjusted p-values are reported in the results unless otherwise stated. Statistical significance was defined as an adjusted p-value < 0.05,

## Acknowledgements

This work was mainly supported by a grant from The Swedish-Norwegian Foundation for Equine Research: grant #H-22-47-717, and in part by FORMAS #2023-01377 and SciLifeLab. The sequencing data were generated with assistance from the SciLifeLab National Genomics Infrastructure, SNP&SEQ Technology Platform, which is funded by the Swedish Research Council and the Knut and Alice Wallenberg Foundation. The raw bulk RNA-Seq analysis process was performed on resources provided by the Uppsala Multidisciplinary Center for Advanced Computational Science (UPPMAX) at Uppsala University and the National Academic Infrastructure for Supercomputing in Sweden (NAISS). We thank all horse owners for participating in the study and the staff at Wången for assistance.

## Author contributions

BZ analysed the data, interpreted results and wrote the manuscript together with AR. HA assisted in sampling, performed the lab work and interpretation of results. MR performed sampling of clinical cases and clinical assessments. MR and SH performed sampling of controls. MH and SH provided expertise in equine asthma. AR conceived and supervised the study, interpreted results, acquired funding and drafted the manuscript. All authors have read and edited the manuscript.

## Data availability

All RNA-sequencing raw data and unprocessed RNA-seq count matrices and metadata analysed in this study will be made publicly available through GEO database.

## Code availability

The code for analysis and the figures in this study will be found on in the github repository: https://github.com/Molmed/Equine_cfRNA.

## Supplementary figure legends

**Supplementary Figure 1.**
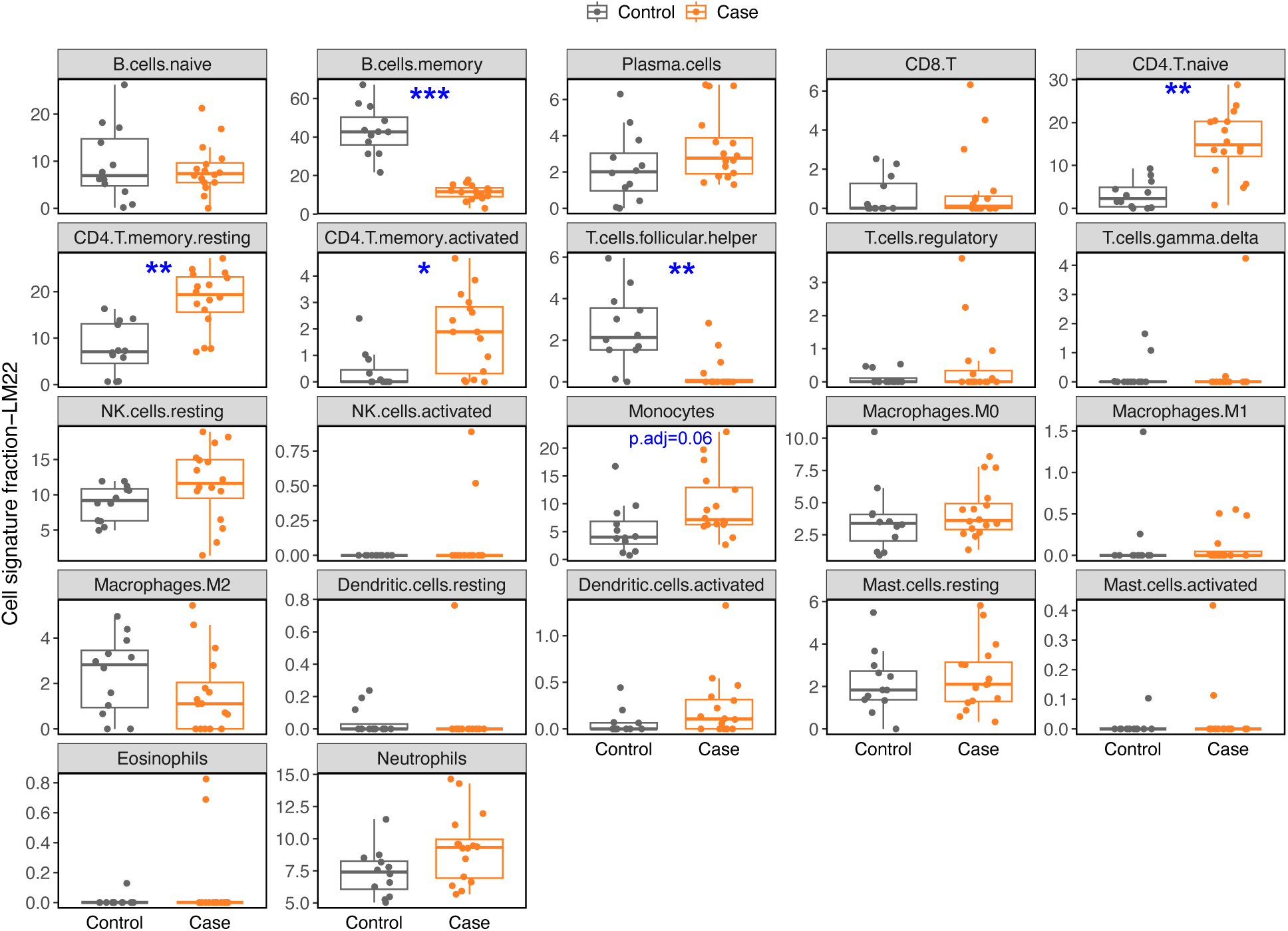
cfRNA cell deconvolution profiling. **A)** Box plots depict relative proportions of 22 distinct immune cell populations estimated using LM22-based deconvolution across control (gray, n = 12) and asthmatic case (orange, n = 16) samples derived from cfRNA sampling. Each panel corresponds to a specific immune cell subset, with cell signature fraction on the y-axis.

**Supplementary Figure 2.**
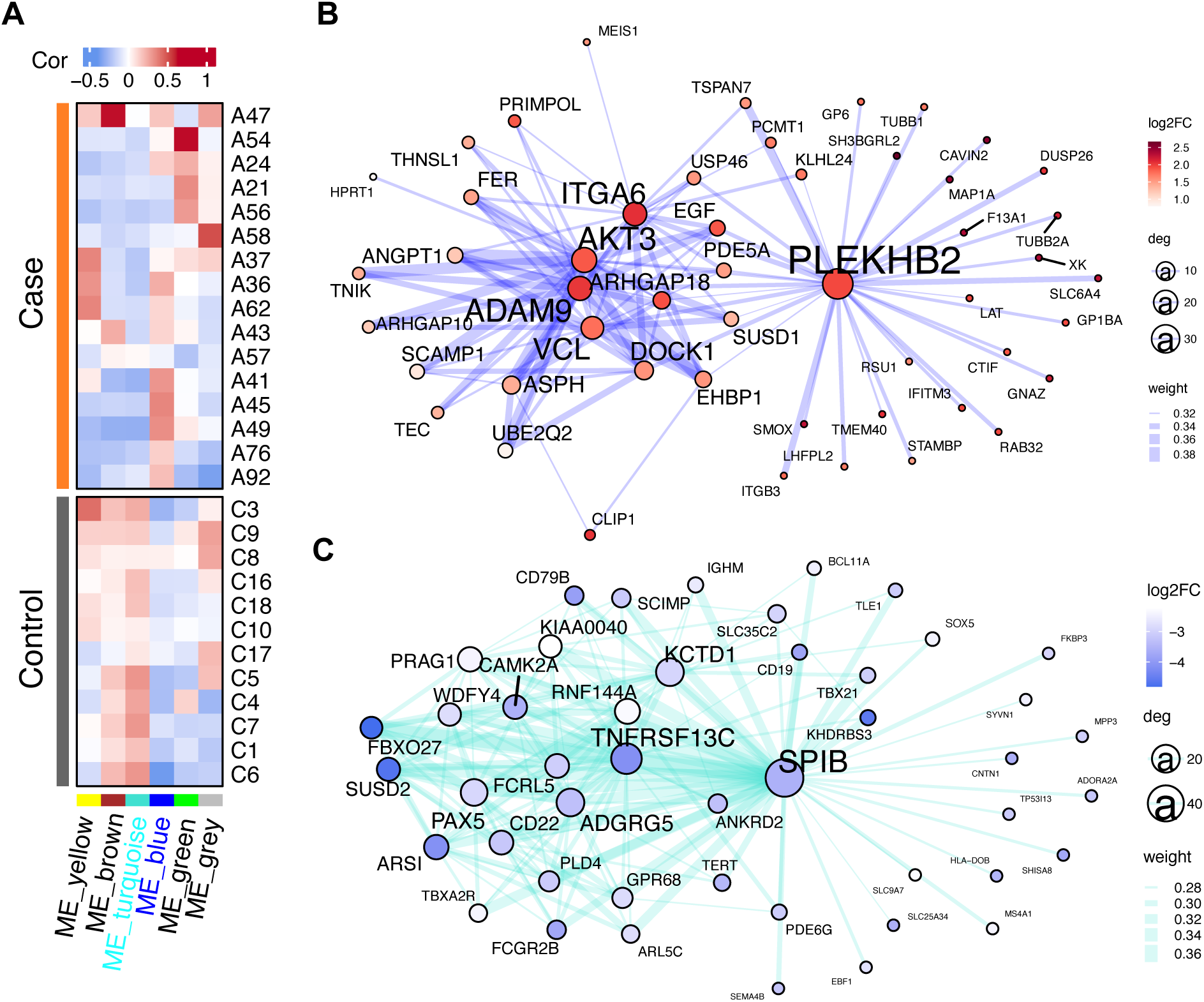
cfRNA WGCNA profiling. (**A**) Weighted gene co-expression network analysis (WGCNA) identified six distinct gene co-expression modules across equine asthma cases (n=16) and controls (n=12). The heatmap displays module eigengene expression patterns across individual samples, with color intensity representing the correlation strength. The blue and turquoise modules showed the significant associations with disease status. (**B, C**) Gene co-expression networks for the blue (B) and turquoise (**C**) modules, illustrating intramodular connectivity and gene–gene correlation patterns. Node color represents log₂ fold change (disease vs. control) from cfRNA differential expression analysis. Node size and gene name font size reflect network connectivity (degree), highlighting hub genes with the most connections. Edge thickness indicates the strength of pairwise gene correlations.

**Supplementary Figure 3.**
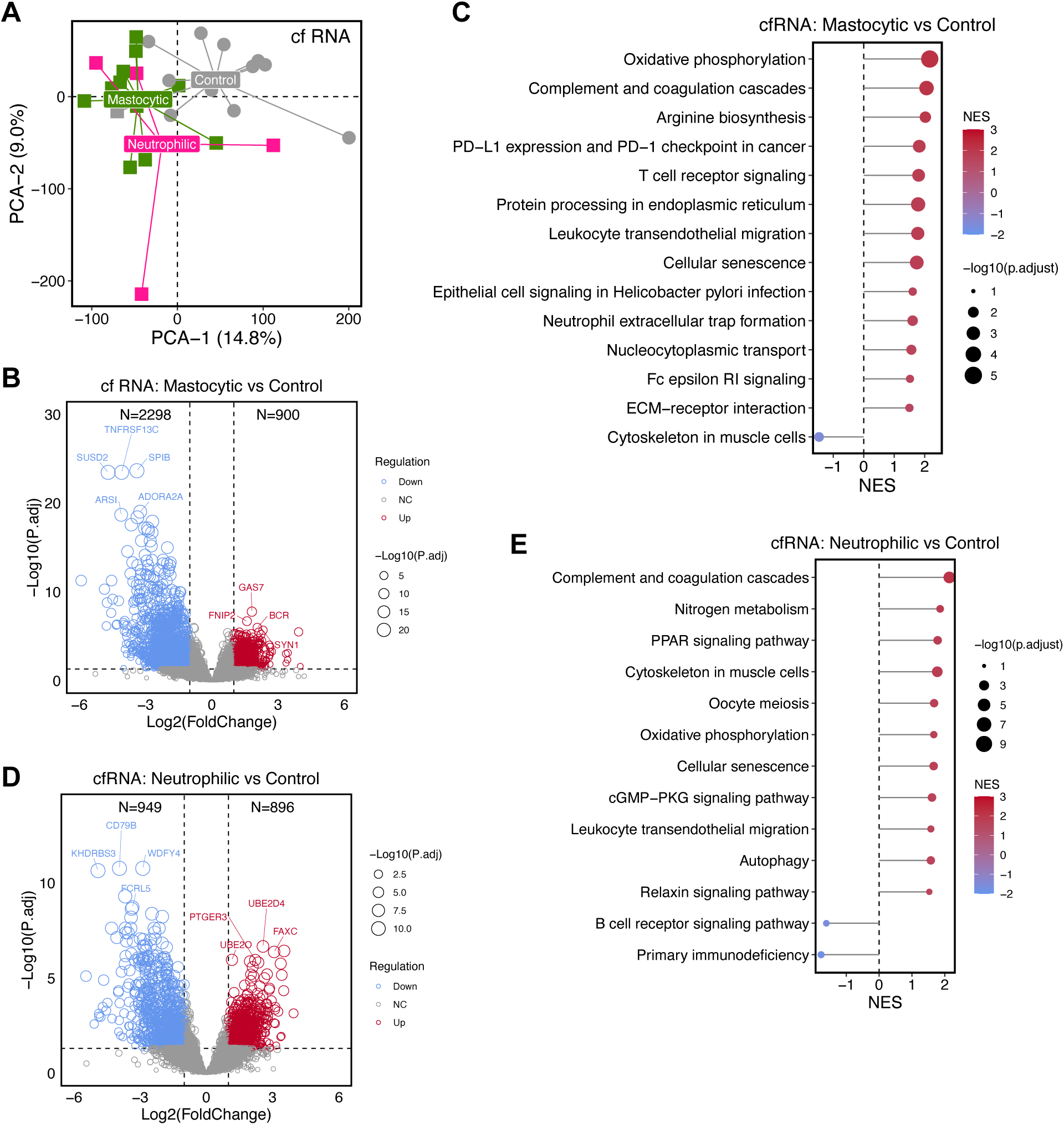
cfRNA profiling based on BAL cytological phenotypes. **A)** PCA of cfRNA gene expression profiles showing separation among control (gray), mastocytic (pink), and neutrophilic (green) phenotypes along PC1 (14.8%) and PC2 (9.0%). **B)** Differential gene expression profile comparing mastocytic and control cfRNA samples. **C)** Differential gene expression profile comparing neutrophilic and control cfRNA samples. **D)** KEGG pathway-level differences between mastocytic and control groups. **E)** KEGG pathway-level differences between neutrophilic and control groups.

**Supplementary Figure 4.**
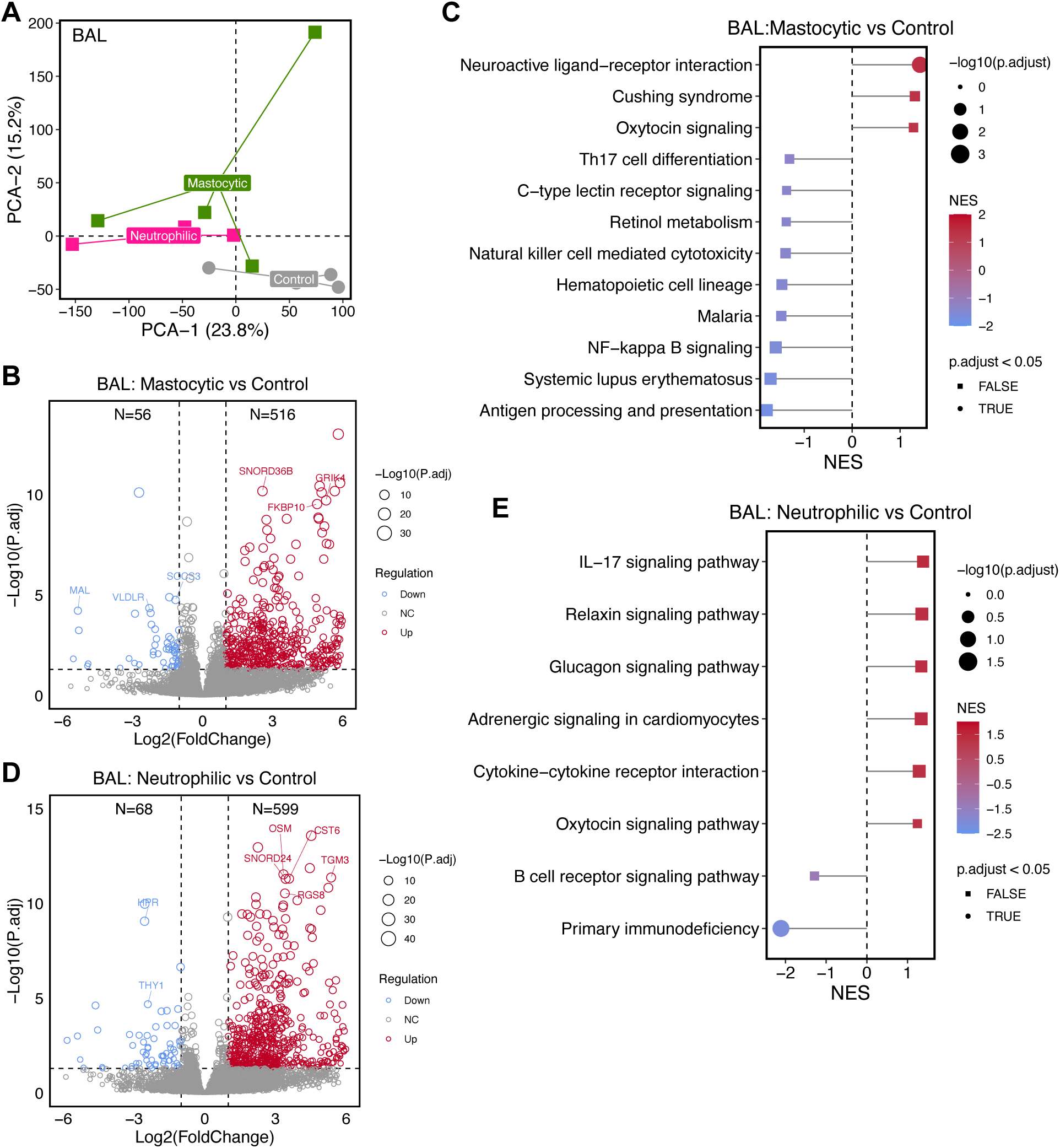
BAL transcriptomic profiling based on cytological phenotypes. **A)** PCA of BAL gene expression profiles showing clear separation among control (gray), mastocytic (pink), and neutrophilic (green) phenotypes along PC1 (14.8%) and PC2 (9.0%). **B)** Differential gene expression profile comparing mastocytic and control BAL samples. **C)** Differential gene expression profile comparing neutrophilic and control BAL samples. **D)** Pathway-level differences between mastocytic and control groups. The majority of pathways were not significant after multiple testing correction. **E)** Pathway-level differences between neutrophilic and control groups. The majority of pathways were not significant after multiple testing correction.

**Supplementary Figure 5.**
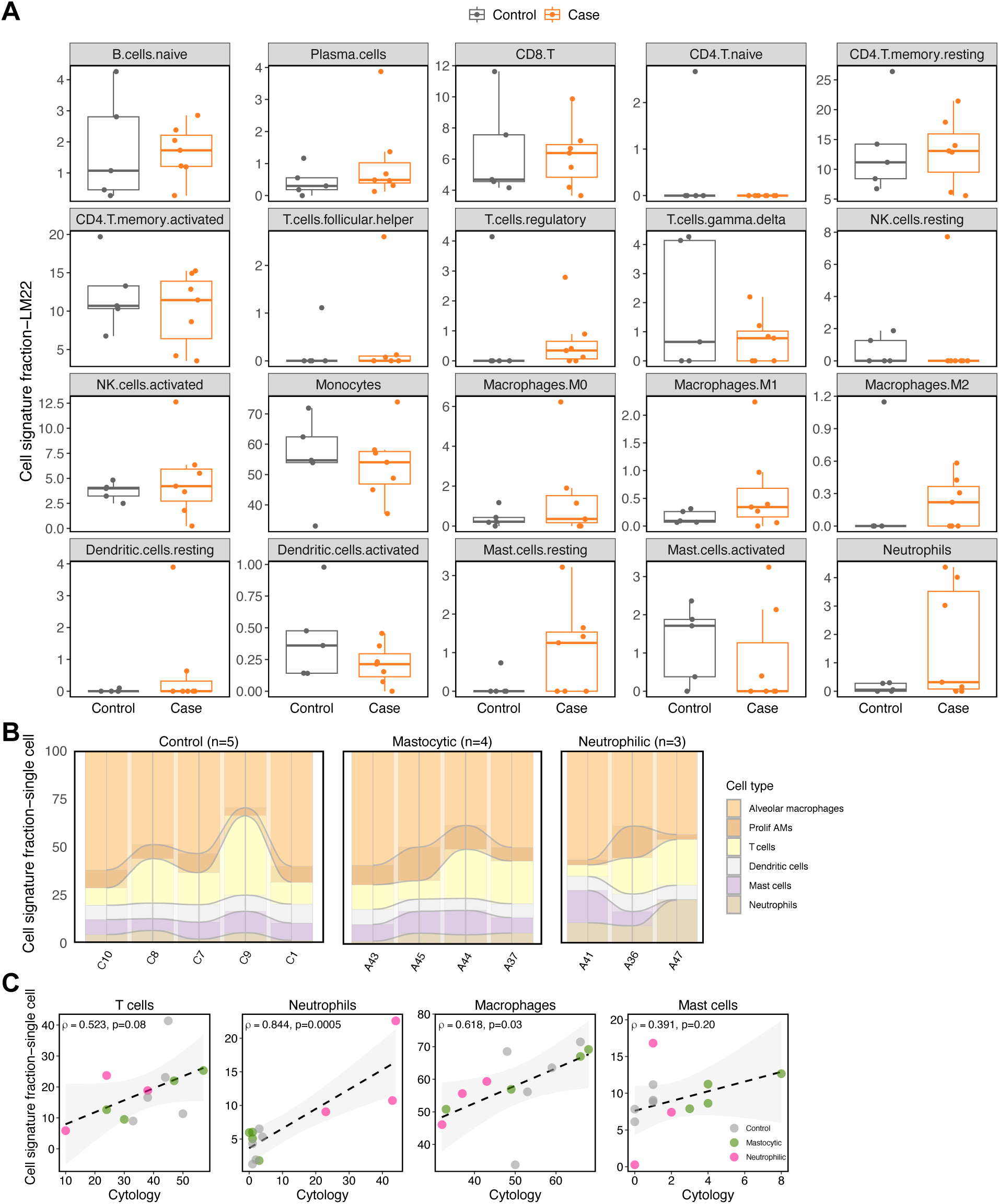
BAL cell deconvolution profiling. **A)** Box plots depict relative proportions of 20 immune cell populations estimated using LM22-based deconvolution across control (gray, n = 5) and asthmatic case (orange, n = 7) samples derived from BAL sampling. Each panel corresponds to a specific immune cell subset, with relative fraction on the y-axis. **B)** Stacked bar chart showing the relative proportions of six cell types across 12 BAL samples, estimated using single-cell reference-based deconvolution. **C)** Validation of single-cell reference-based computational deconvolution. Scatter plots correlate computationally inferred cell fractions (y-axis) with cytological measurements (x-axis) for four cell types. Each point represents an individual BAL sample, colored by cytological phenotype (gray: control; pink: mastocytic; green: neutrophilic).

**Supplementary Figure 6.**
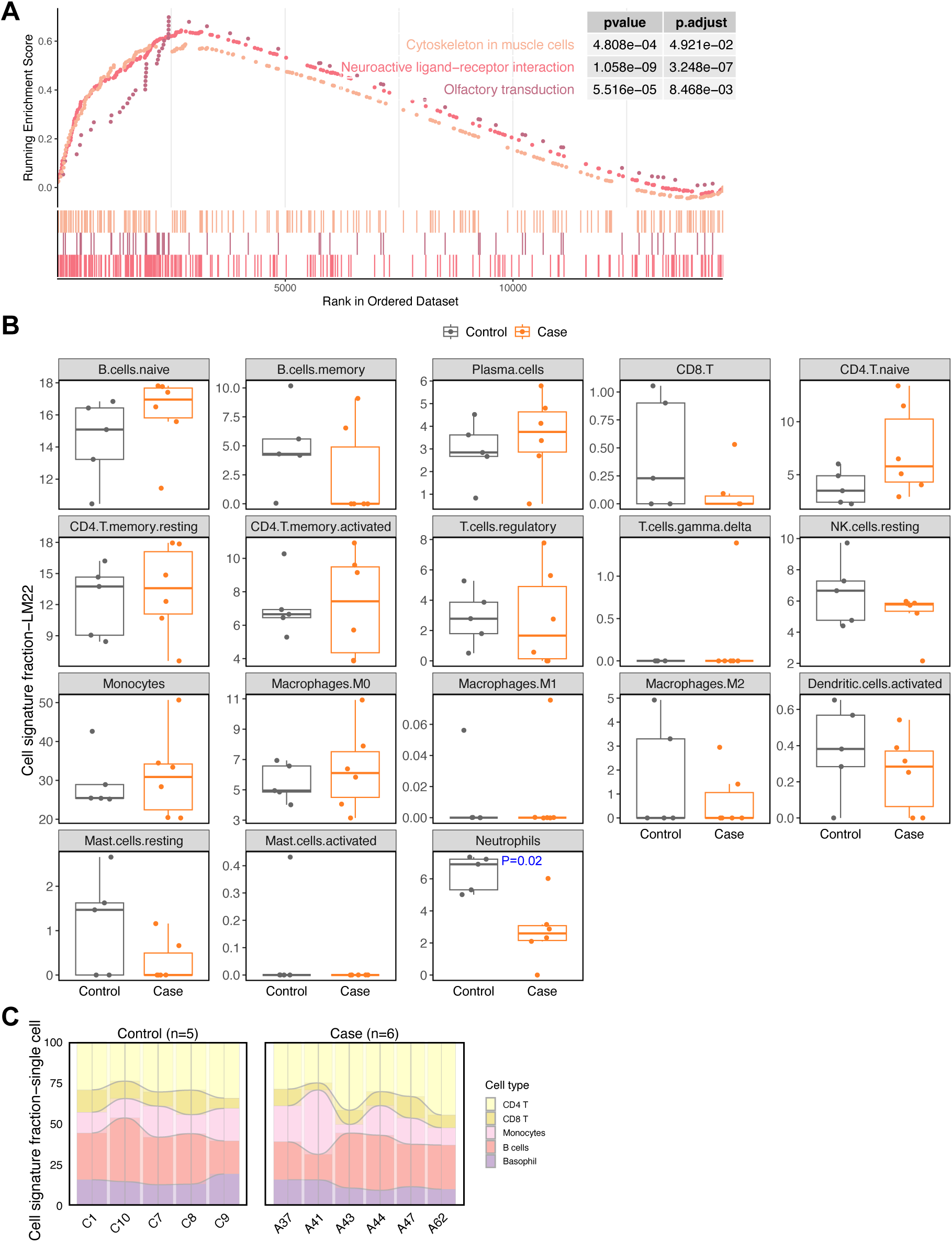
Whole blood transcriptomic and cellular profiling. **A)** Gene Set Enrichment Analysis (GSEA). Representative enrichment plots for three significantly altered pathways are shown; peak enrichment scores and gene positions highlight pathways significantly activated in the case group. **B)** Stacked bar chart depicting average relative proportions of five immune cell types estimated by single-cell reference-based deconvolution across samples. **C)** Box plots show relative proportions of 18 distinct immune cell populations estimated using LM22-based deconvolution across control (gray, n = 5) and asthmatic case (orange, n = 6) samples derived from WB sampling. Each panel represents a specific immune cell subset, with relative fraction on the y-axis.

