## Supplementary Table 1 for "Circulating cell-free RNA reflects inflammatory and airway remodeling signatures in an equine model of asthma"

| ID | Macrophage | T cells | Neutrophil | Eosinophil | Mast cells | Phenotype | Class | Comp^#^ | Age | Sex | Breed | Ongoing CS treatments | Other |
| --- | --- | --- | --- | --- | --- | --- | --- | --- | --- | --- | --- | --- | --- |
| A92 | 54 | 38 | 3 | 1 | 4 | mastocytic | mEA | cf | 16 | mare | Warmblood | No |  |
| A47 | 32 | 24 | 44 | 0 | 0 | neutrophilic | sEA | cf/BAL/WB | 20 | gelding | Icelandic | No | ASIT |
| A62 | 77 | 9 | 6 | 1 | 7 | mastocytic | mEA | cf/WB | 4 | gelding | Arabian horse | No |  |
| A56 | 59 | 37 | 1 | 1 | 2-4* | mastocytic | mEA | cf | 19 | gelding | Icelandic | No |  |
| A57 | 56 | 39 | 2 | 0 | 3 | mastocytic | mEA | cf | 14 | mare | Warmblood | No | ASIT |
| A43 | 68 | 24 | 3 | 1 | 4 | mastocytic | mEA | cf/BAL/WB | 14 | mare | Shetland pony | No |  |
| A37 | 49 | 47 | 1 | 0 | 3 | mastocytic | mEA | cf/BAL/WB | 12 | gelding | Warmblood | Two weeks prior BAL | ASIT, Seasonal EA |
| A49 | 66 | 21 | 1 | 0 | 11 | mastocytic | mEA | cf | 7 | mare | Shetland pony | No |  |
| A36 | 37 | 38 | 23 | 0 | 2 | neutrophilic | sEA | cf/BAL | 18 | gelding | Warmblood | No |  |
| A24 | 71 | 24 | 4 | 0 | 5 | mastocytic | mEA | cf | 11 | gelding | Icelandic | No | ASIT, Seasonal EA |
| A54 | 58 | 30 | 2 | 0 | 10 | mastocytic | mEA | cf | 18 | gelding | Connemara | No |  |
| A58 | 53 | 29 | 4 | 0 | 14 | mastocytic | mEA | cf | 8 | gelding | Welshpony | No |  |
| A45 | 66 | 30 | 0 | 0 | 4 | mastocytic | mEA | cf/BAL | 11 | gelding | New Forest | No |  |
| A76 | 69 | 29 | 0 | 0 | 2 | paucigranulocytic | mEA | cf | 5 | gelding | Warmblood | No | coughing & excercise intolerant, |
| A44 | 33 | 57 | 1 | 1 | 8 | mastocytic | mEA | BAL/WB | 14 | mare | Warmblood | No |  |
| A41 | 43 | 10 | 43 | 0 | 1 | neutrophilic | sEA | cf/BAL/WB | 15 | mare | Warmblood | Yes |  |
| A21 | 48 | 36 | 15 | 0 | 1 | neutrophilic | sEA^1^ | cf | 19 | gelding | Warmblood | Yes |  |
| C1 | 48 | 50 | 1 | 0 | 1 | control |  | cf/BAL/WB | 5 | gelding | Standardbred | No |  |
| C3 | 46 | 52 | 2 | 0 | 0 | control |  | cf | 10 | gelding | Standardbred | No |  |
| C4 | 53 | 43 | 1 | 1 | 1 | control |  | cf | 11 | gelding | Standardbred | No |  |
| C5 | 42 | 50 | 6 | 0 | 2 | control |  | cf | 11 | mare | Standardbred | No |  |
| C6 | 39 | 51 | 7 | 0 | 3 | control |  | cf | 8 | mare | Standardbred | No |  |
| C7 | 59 | 38 | 2 | 0 | 1 | control |  | cf/BAL/WB | 10 | gelding | Standardbred | No |  |
| C9 | 50 | 45 | 4 | 0 | 1 | control |  | cf/BAL/WB | 11 | gelding | Standardbred | No |  |
| C10 | 66 | 33 | 1 | 0 | 0 | control |  | cfBAL/WB | 6 | gelding | Standardbred | No |  |
| C16 | 39 | 60 | 0 | 0 | 1 | control |  | cf | 13 | gelding | Standardbred | No |  |
| C17 | 41 | 57 | 2 | 0 | 0 | control |  | cf | 6 | mare | Standardbred | No |  |
| C18 | 71 | 27 | 0 | 0 | 2 | control |  | cf | 6 | mare | Standardbred | No |  |
| C8 | 53 | 44 | 3 | 0 | 0 | control |  | cf/BAL/WB | 7 | gelding | Standardbred | No |  |

*varied among different slide preparations

\# Compartments sampled

1. This horse previously have had 50 % neutrophils in BAL

**Supplementary Table 1. BAL cytology and phenotypes of cases & controls**
