## Supplementary Table 2 for "Circulating cell-free RNA reflects inflammatory and airway remodeling signatures in an equine model of asthma"

***cfRNA DGE signatures: Functional themes, Representative Genes (Up / Down), Interpretation***

Relevant DEG genes were grouped into modules according to their function (as they are reported in the literature)

**Inflammation & Remodeling Mediators**

Up: TGFB2, MMP1, MMP8, HPSE, APP, SERPINE1, ADAM9, TIMP4

Down: TGFB3, WNT5A, WNT7A, ERRFI1, MMP14, MMP28, COL1A1, ADAMTS14/17

Interpretation: Up: injury/inflammation + ECM breakdown. Down: new collagen/structural assembly

**Epithelial & Barrier-Associated Genes**,

Up: IL33, EGF, FGF1, ECM1, KRT16

Down: SFTPD, CLDN3/9/14, KRT18/12/40, CDH13/20/26

Interpretation: Shift toward epithelial stress/alarmin release, barrier genes down

**Chemokines, Cytokines & Receptors**

Up: CCR4, CXCR4, IL7R, IL13RA1, IL12RB2, EGF, FGF1

Down: IL11. CSF3, TNFSF15, PGF, EDN2, STC1, IL12B, IL21R, IL20RA, IFNGR1, IFNLR1, TNFRSF11A/19/9, CCL24, CCL26, CXCR5, CXCR1

Intepretation Tilt toward Th2 + tissue remodeling; Th1/IFN/eotaxin programs down

**Mast Cell & Granulocyte-Related**

Up: IGHE, SRGN, PTGS1

Down: FCER2, CCL24, CCL26, CSF3, CYBB, CLEC5A, SIGIRR, PADI2, TBXA2R

Interpretation: IgE–mast cell axis up, eosinophil chemoattractants and myeloid regulators down

**Eicosanoid/Leukotriene Pathway**

Up: ALOX5AP, PTGS1

Down: PLA2G6, PLA2G4F, TBXA2R

Interpretation: Mixed, Leukotriene/prostanoid potential up; thromboxane signaling down

**Immune Signaling & Activation**,

Up: IL7R, IL13RA1, CD40LG, CD28, CD4, CD69, LAT, ITK, CCR4, CXCR4, GRAP2, NFIL3

Down: CD19, CD79A/B, MS4A1, CD22, BANK1, BLK, POU2AF1, PAX5, EBF1, SPIB, FCRL1/3/4/5, TNFRSF13B/C, CD72, CTLA4, CD40, CIITA, CD74, HLA-DOA/B, SYK, IRAK1/2, TRAF3, TRIM25, INPP5D, NFKB2, RELB, NFKBIE

Interpretation: Clear T-cell activation up, B-cell/APC/programs down

**Platelet/Endothelial Activation**

Up: VWF, SELP, PECAM1, ITGA2/6/1/2B, ITGB1/3, ITGAX

Down: SELE, PGF, EFNA1, ROBO1, ROBO2

Interpretation: Platelet adhesion/activation up, endothelial adhesion genes variable

**Interferon-Stimulated Genes (ISGs**)

Up: STAT1, ISG20, IRF2, CD226

Down: IFNLR1, IFNGR1, IRF3, IRF5, IFI30, CIITA, HLA-DOA/B

Interpretation: STAT1 stress signaling up, canonical IFN receptor / IRF and some APC down
